# The Arabidopsis NRT1/PTR FAMILY Protein NPF7.3/NRT1.5 is an Indole-3-butyric Acid Transporter Involved in Root Gravitropism

**DOI:** 10.1101/2020.06.04.131797

**Authors:** Shunsuke Watanabe, Naoki Takahashi, Yuri Kanno, Hiromi Suzuki, Yuki Aoi, Noriko Takeda-Kamiya, Kiminori Toyooka, Hiroyuki Kasahara, Ken-Ichiro Hayashi, Masaaki Umeda, Mitsunori Seo

## Abstract

Active membrane transport of plant hormones and their related compounds is an essential process that determines the distribution of the compounds within plant tissues and, hence, regulates various physiological events. Here, we report that the Arabidopsis NITRATE TRANSPORTER 1/PEPTIDE TRANSPORTER FAMILY 7.3 (NPF7.3) protein functions as a transporter of indole-3-butyric acid (IBA), a precursor of the major endogenous auxin indole-3-acetic acid (IAA). When expressed in yeast, NPF7.3 mediated cellular IBA uptake. Loss-of-function *npf7.3* mutants showed defective root gravitropism with reduced IBA levels and auxin responses. Nevertheless, the phenotype was restored by exogenous application of IAA but not by IBA treatment. *NPF7.3* was expressed in pericycle cells and the root tip region including root cap cells of primary roots where the IBA-to-IAA conversion occurs. Our findings indicate that NPF7.3-mediated IBA uptake into specific cells is required for the generation of appropriate auxin gradients within root tissues.

---

Plant hormones are vital compounds that function as signaling molecules and control a wide range of physiological responses so that plants can cope with fluctuating environments^1^. Often plant hormones induce physiological responses very locally^2-7^; thus, the endogenous levels as well as the signal transduction pathways of plant hormones are regulated very precisely. Endogenous concentrations of bioactive plant hormones are primarily determined by the balance between the rates of biosynthesis and degradation (or inactivation)^1^. Additionally, specific transport systems for plant hormones and their related compounds are also considered to be critical components for generating appropriate hormone concentrations within cells, since hormone biosynthesis and perception do not necessarily take place in the same cell. In fact, movement of plant hormones and their precursors has been reported^8^.

Auxin is the best-characterized mobile plant hormone that plays crucial roles in many aspects of plant growth and development by mainly regulating cell division, cell elongation and cell differentiation^1^. Indole-3-acetic acid (IAA) is the major naturally occurring auxin. Measurements of IAA by mass spectrometry as well as visualization of auxin responses by reporter genes have suggested that IAA is differentially distributed within plant tissues. Furthermore, appropriate IAA gradients or local IAA maxima/minima are required for proper auxin functions. In fact, mutants impaired in IAA transport have been isolated based on their defects in organ development and various physiological responses. Evidence for IAA transport in a cell-to-cell manner by combinations of several transmembrane transporters is well documented. The plant-specific PIN-FORMED (PIN) transporters and the AUX1/LAX family of amino acid permease-like proteins are IAA efflux and influx carriers, respectively^9,10^. PINs determine the direction of IAA flow by localizing unevenly in the plasma membrane, whereas polarity is not observed for AUX/LAXs distribution. Also, some subfamily B members of the ATP-binding cassette (ABC) transporter (ABCBs) family, act as IAA transporters^11^.

IAA is mainly synthesized from tryptophan via indole-3-pyruvic acid, however, minor routes for IAA biosynthesis have also been suggested. Indole-3-butyric acid (IBA) has auxin activity when applied exogenously to plants. Arabidopsis mutants isolated based on their resistance to exogenous IBA are defective in the conversion of IBA to IAA. Since IBA has been detected in several plant species, the compound is considered to be an endogenous IAA precursor. Possible IBA transporters have been identified^12-14^, suggesting that IBA transport is also an important factor that regulates endogenous IAA levels.

NITRATE TRANSPORTER 1/PEPTIDE TRANSPORTER FAMILY (NPF) proteins have been characterized as transporters for several plant hormones including auxin, although the family was initially identified as nitrate or di/tri-peptide transporters^15,16^. The Arabidopsis NPF6.3 protein (also called as CHL1 or NRT1.1) is a well-characterized dual-affinity nitrate transporter that mediates nitrogen uptake from the rhizosphere, however, the same protein also transports IAA^17,18^. More recently, the Arabidopsis NPF5.12 protein, also named TRANSPORTER OF IBA1 (TOB1), was identified as an IBA transporter that localizes to the vacuolar membrane^14^. Here, we report that NPF7.3, originally identified as a low-affinity nitrate transporter (NRT1.5) and later shown to be a proton/potassium antiporter in Arabidopsis^19,20^, functions as an IBA transporter. We found that *npf7.3* mutant roots were not able to respond properly to gravity. Our transport assays in yeast demonstrated that NPF7.3 efficiently mediated IBA uptake into the cells. Asymmetric distribution of auxin activities in root tips following the reorientation of roots was not observed in the *npf7.3* mutants, and the defect was restored by IAA but not by IBA treatment. These results suggest that cellular IBA uptake mediated by NPF7.3 contributes to the production of IAA required to fully induce Arabidopsis root gravitropism.

## Results

### Mutants defective in NPF7.3 exhibit altered root gravitropism

Some of the 53 members of the Arabidopsis NPF proteins transport plant hormones such as IAA, abscisic acid (ABA), gibberellin (GA) and jasmonates (jasmonoyl isoleucine; JA-Ile)^16^. Thus, we hypothesized that there are still unidentified plant hormone transporters in this protein family. We examined the physiological functions of several members of the NPF proteins more closely based on their transport activities in yeast and/or their expression patterns available from public databases. NPF7.3 was one of the candidates since its gene expression was affected by biotic and abiotic stresses as well as IAA treatment (eFP Browser, http://bar.utoronto.ca/efp/cgi-bin/efpWeb.cgi^21^). NPF7.3 was originally characterized as a low-affinity bidirectional nitrate transporter NRT1.5 that facilitates xylem loading of nitrate from root pericycle cells^19^. NPF7.3 has also been reported to function as a proton/potassium antiporter (also referred to as LKS2) that mediates xylem loading of potassium^20^. Previous studies showed that mutants defective in NPF7.3 exhibit phenotypes related to root growth especially when nitrate or potassium concentrations in the growth media were low; however, we found that two allelic mutants of NPF7.3 showed a wavy root growth phenotype when grown on vertical plates containing half strength Murashige-Skoog (MS) medium, which is rich in nitrate and potassium (more than 10 mM ammonium nitrate and 9 mM potassium nitrate) and 1% sucrose (Fig. 1a). In our study, we used a previously characterized mutant allele (*nrt1.5-4*) and a newly isolated allele (Supplementary Fig. 1), and tentatively named them *npf7.3-4* and *npf7.3-6* (five *nrt1.5* alleles have been reported so far), respectively, for simplification. As compared with wild type, *npf7.3* mutant roots had significantly increased wave frequencies with reduced wave intervals, whereas primary root lengths were almost comparable in the wild type and *npf7.3* (Fig. 1b-d). Consequentially, *npf7.3* mutants had lower root straightness indices^22^ compared to wild type (Fig. 1e). Since several Arabidopsis mutants with wavy root growth display altered gravitropism^23,24^, we investigated the phenotype in *npf7.3* primary roots. Indeed, gravitropic indices^22^ in the *npf7.3* mutants were significantly lower than those in wild type (Fig. 1f). To further validate the defects of *npf7.3* in root gravitropic responses, vertical plates containing wild-type and *npf7.3* seedlings were rotated to shift the gravity direction 90° (1^st^ rotation), and then the plates were rotated again back to the initial state (2^nd^ rotation) after 24 hours of incubation. This experiment was conducted in the absence of sucrose in the media, since the wavy root phenotype interferes with the measurement of root angles. Although most wild-type roots properly change their growth direction toward gravity (nearly 90°), *npf7.3* mutant roots responded poorly to gravity (Fig. 1g). As a result, root angles were significantly larger and variable in *npf7.3* compared with wild type; however, root elongation rates after reorientation of plants were comparable between wild type and the mutant (Fig. 1h, i). We sometimes observed that root tips of *npf7.3* were not attached to the surface of agar plates after reorientation. These plants were not used for measuring root angles and root growth rates. Finally, we confirmed that the disruption of root gravitropism observed in *npf7.3* was restored by the expression of wild-type *NPF7.3* ORF under the control of its own promoter (2.1 kb) (Fig. 1j).

**Fig. 1.**
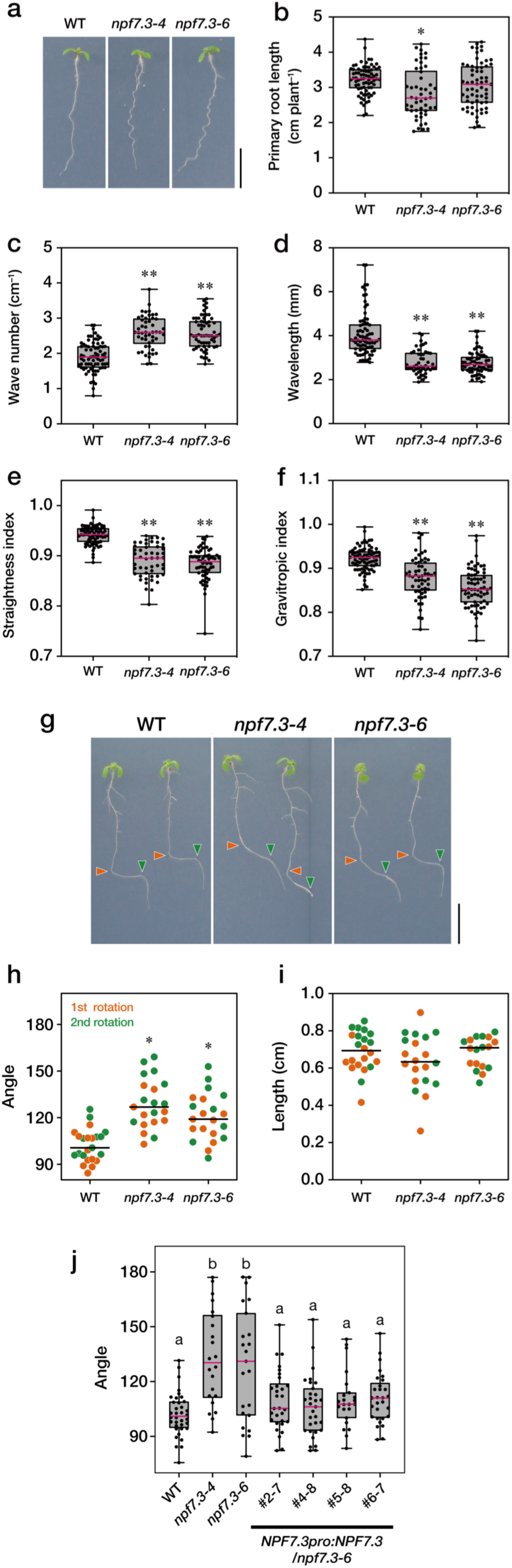
Wavy root growth and altered root gravitropism observed in *npf7.3*. **a**. Representative images of seven-day-old wild-type (WT) and *npf7.3* (*npf7.3-4* and *npf7.3-6*) seedlings grown on half-strength MS media containing 1.5% (w/v) agar and 1% (w/v) sucrose. Scale bar = 1 cm. **b-f**. Lengths (**b**), wave numbers (**c**), wave lengths (**d**), straightness indices (**e**), and gravitropic indices (**f**) of primary roots measured in seven-day-old wild-type (WT) and *npf7.3* (*npf7.3-4* and *npf7.3-6*) seedlings. The straightness index is defined as the ratio of the direct distance from the origin of the root to the root tip and the primary root length. The gravitropic index is defined as the ratio of the root tip ordinate and the primary root length. Dots represent individual measurements, horizontal pink bars represent the median, boxes represent the middle 50% of the distribution, and whiskers represent the entire spread of the data. Asterisks indicate statistically significant differences compared to wild-type determined by Tukey’s multiple comparison test (*P* < 0.05). **g**. Representative images of wild-type (WT) and *npf7.3* (*npf7.3-4* and *npf7.3-6*) seedlings after reorientation. Vertical plates containing seven-day-old seedlings were rotated 90°, and the plates were again rotated 90° back to the initial position after 24 hours. Photos were taken 24 hours after the second rotation. Orange and green triangles indicate the positions of root tips at the time of the first and second rotations, respectively. Scale bar = 1 cm. **h,i**. Angles and lengths of primary roots 24 hours after the first and second rotations. Dots indicate individual measurements. Lines represent the medians of each group. Asterisks indicate significant differences compared with wild type by Dunn’s multiple comparison test (*P* < 0.001). **j**. Complementation of *npf7.3* with wild-type *NPF7.3* CDS. Root angles of wild type (WT), *npf7.3* (*npf7.3-4* and *npf7.3-6*) and *npf7.3* expressing wild-type NPF7.3 (*NPF7.3pro:NPF7.3*/*npf7.3*) were measured 24 hours after the first rotation. Horizontal pink bars represent the median, boxes represent the middle 50% of the distribution, and whiskers represent the entire spread of the data. Different letters indicate statistically significant differences by Dunn’s multiple comparison test (*P* < 0.05).

### NPF7.3 has IBA uptake activity

The plant hormone auxin plays a crucial role in root gravitropism. Therefore, we hypothesized that NPF7.3 somehow regulates auxin responses. Given that NPFs transport a variety of compounds including IAA, it is possible that NPF7.3 transports IAA and/or its related compounds. To assess this possibility, we conducted direct transport assays using yeast cells. Significantly larger amounts of IAA were accumulated in NPF7.3-expressing cells compared to the cells containing a control vector when the chemical concentration was 10 µM and the reaction buffer was acidic (pH 5.8) (Fig. 2a). We also examined whether IBA could be a substrate for NPF7.3 in the same conditions because the compound is considered to be an IAA precursor^25,26^. Interestingly, IBA also accumulated at a much higher level in the cells expressing NPF7.3 compared to the control cells (Fig. 2b). Moreover, IBA was more efficiently taken into the cells expressing NPF7.3 than IAA.

**Fig. 2.**
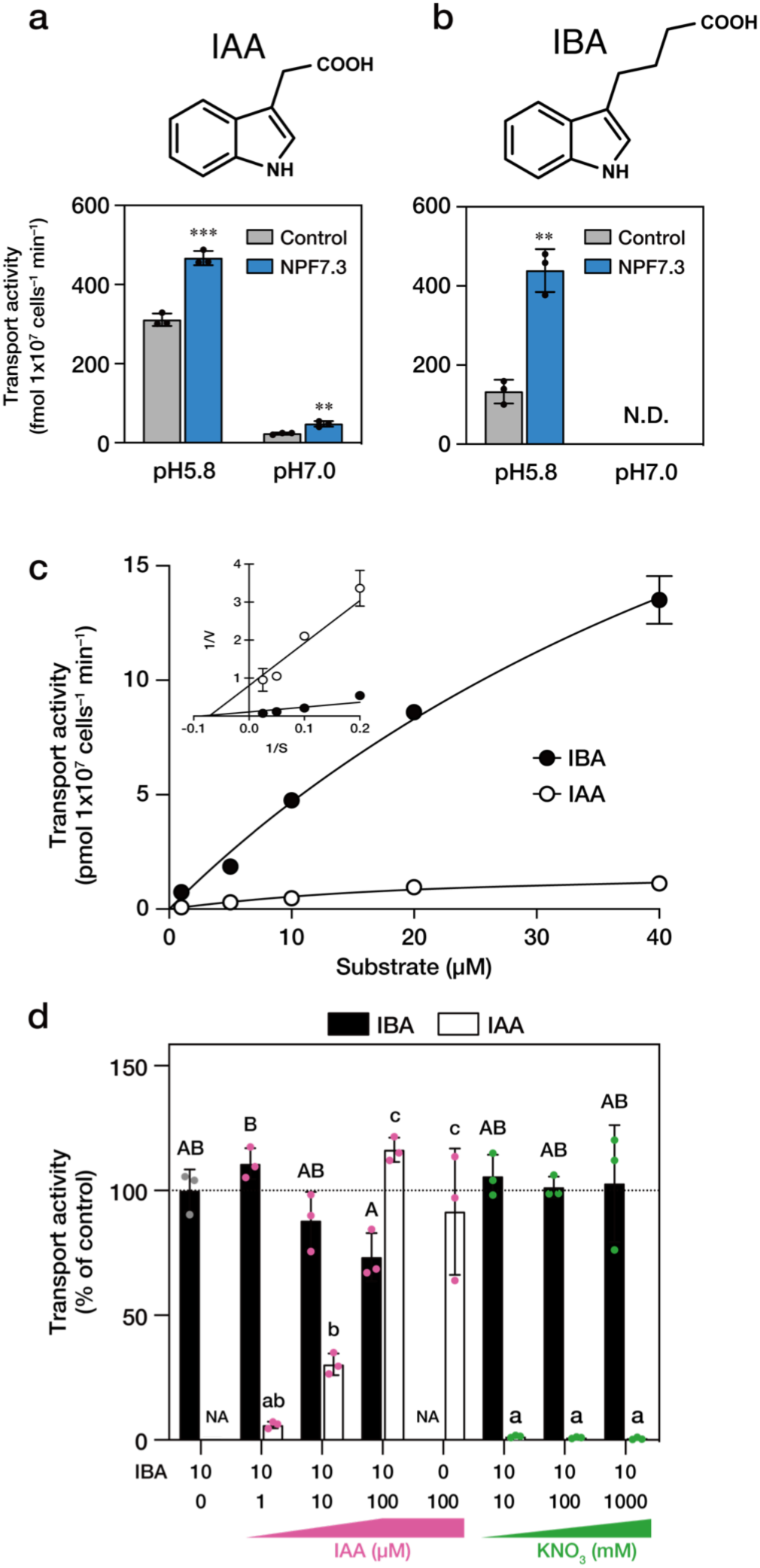
IAA and IBA transport activities of NPF7.3. **a,b**. Activities of NPF7.3 to transport IAA and IBA into yeast cells when the substrate concentrations were 10 µM. Yeast cells that were transformed with the empty pYES2-DEST52 vector were used as a control. Bars indicate standard deviations of the means. Asterisks indicate statistically significant differences compared to the controls (*P* < 0.001 by Student’s *t*-test). N.D., not detected. **c**. Dose dependent uptake of IBA and IAA mediated by NPF7.3 at pH5.8. The amount of IBA or IAA taken into the yeast cells expressing *NPF7.3* were determined when the substrate concentrations were 0, 5, 10, 20, and 40 µM. Data are shown as means ± SD of three independent experiments. The inset shows a Lineweaver–Burk plot. **d**. Effects of nitrate and IAA on NPF7.3-mediated IBA uptake. The amount of IBA taken into the yeast cells expressing *NPF7.3* in the presence of potassium nitrate (KNO_3_) or IAA at indicated concentrations was determined. IAA was also quantified when it was present as a competitor. Data are means ± SD of three independent experiments. Different uppercase and lowercase letters indicate statistically significant differences by one-way ANOVA with Tukey’s multiple comparison test (*P* < 0.05) for IBA and IAA levels, respectively. NA means not analyzed.

Kinetic analyses indicated that the *K*m values of NPF7.3 for IAA and IBA were comparable (14 µM and 11 µM for IAA and IBA, respectively), however, NPF7.3 had a much higher *V*max value for IBA compared to IAA (9 pmols 1×10^7^ cells^−1^ min^−1^ for IBA and 1.2 pmoles 1×10^7^ cells^−1^ min^−1^ for IAA) (Fig. 2c). When both IAA and IBA were present at the same concentration (10 µM) as substrates of NPF7.3, IBA was more preferentially transported into the yeast cells (Fig 2d). Furthermore, IBA uptake mediated by NPF7.3 was only slightly inhibited when 10 times higher concentration of IAA relative to IBA was present as a competitor (100 µM IAA versus 10 µM IBA) (Fig. 2d). These results indicate that IBA is a better substrate for NPF7.3 than IAA. On the other hand, no significant effects on IBA uptake were observed even when the concentration of potassium nitrate (KNO_3_) was 10,000 times higher than that of IBA (100 mM KNO_3_ versus10 µM IBA) in the reaction buffer.

### Mutations in *NPF7.3* reduced endogenous IBA levels

To determine the possible involvement of NPF7.3 in IBA transport *in vivo*, we first measured the endogenous IBA and IAA content in plant tissues. We found that IBA levels in *npf7.3* roots were reduced to approximately one-half of the wild-type levels (Fig. 3), whereas IBA in the shoots of both wild type and *npf7.3* was undetectable. In contrast, IAA levels in both roots and shoots were unchanged in *npf7.3* compared to wild type. We also determined endogenous levels of IAA metabolites in roots, however, these compounds accumulated to similar levels both in wild type and *npf7.3* (Supplementary Fig. 2). In addition, no significant difference was observed for endogenous levels of other plant hormones between wild type and *npf7.3* (Supplementary Fig. 3).

**Fig. 3.**
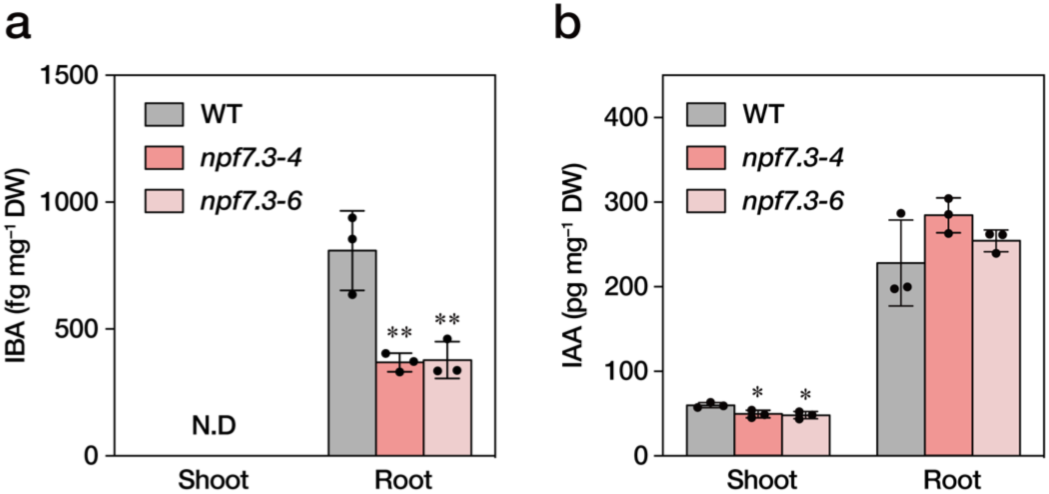
Endogenous levels of IBA and IAA in *npf7.3*. IBA and IAA levels in shoots and roots of 14-day-old wild-type (WT) and *npf7.3* (*npf7.3-4* and *npf7.3-6*) seedlings were quantified. Bars indicate standard deviations of the means. Dots represent individual measurements. Asterisks indicate significant differences compared with wild-type (\**P* < 0.05 by Tukey’s multiple comparison test).

### Exogenous supplementation with IAA but not IBA restored defective gravitropic responses of *npf7.3*

IBA is proposed to be converted to IAA through a process similar to fatty acid ß-oxidation in peroxisomes, at least when applied exogenously to Arabidopsis^27^. Therefore, we hypothesized that IBA taken inside the cells mediated by NPF7.3 has to be converted to IAA to induce physiological responses. To test this hypothesis, we investigated the effects of exogenous IBA and IAA treatments on root gravitropism of *npf7.3*.In control media, the angles of wild-type roots were mostly 90°, whereas *npf7.3* roots responded less to gravity and had larger root angles compared to wild type (approximately 120°). Addition of 10 or 50 nM IAA to the media did not affect the wild-type responses to gravity, however, the defective gravitropic responses observed in *npf7.3* were restored at least partially when 50 nM IAA was present (Fig. 4). IBA supplementation (10 to 200 nM) slightly increased the population of wild-type roots that responded less to gravity, but the majority of roots were still bent at a 90° angle. In these conditions, in contrast to the case with IAA treatment, IBA did not improve the impaired gravitropism of *npf7.3*.

**Fig. 4.**
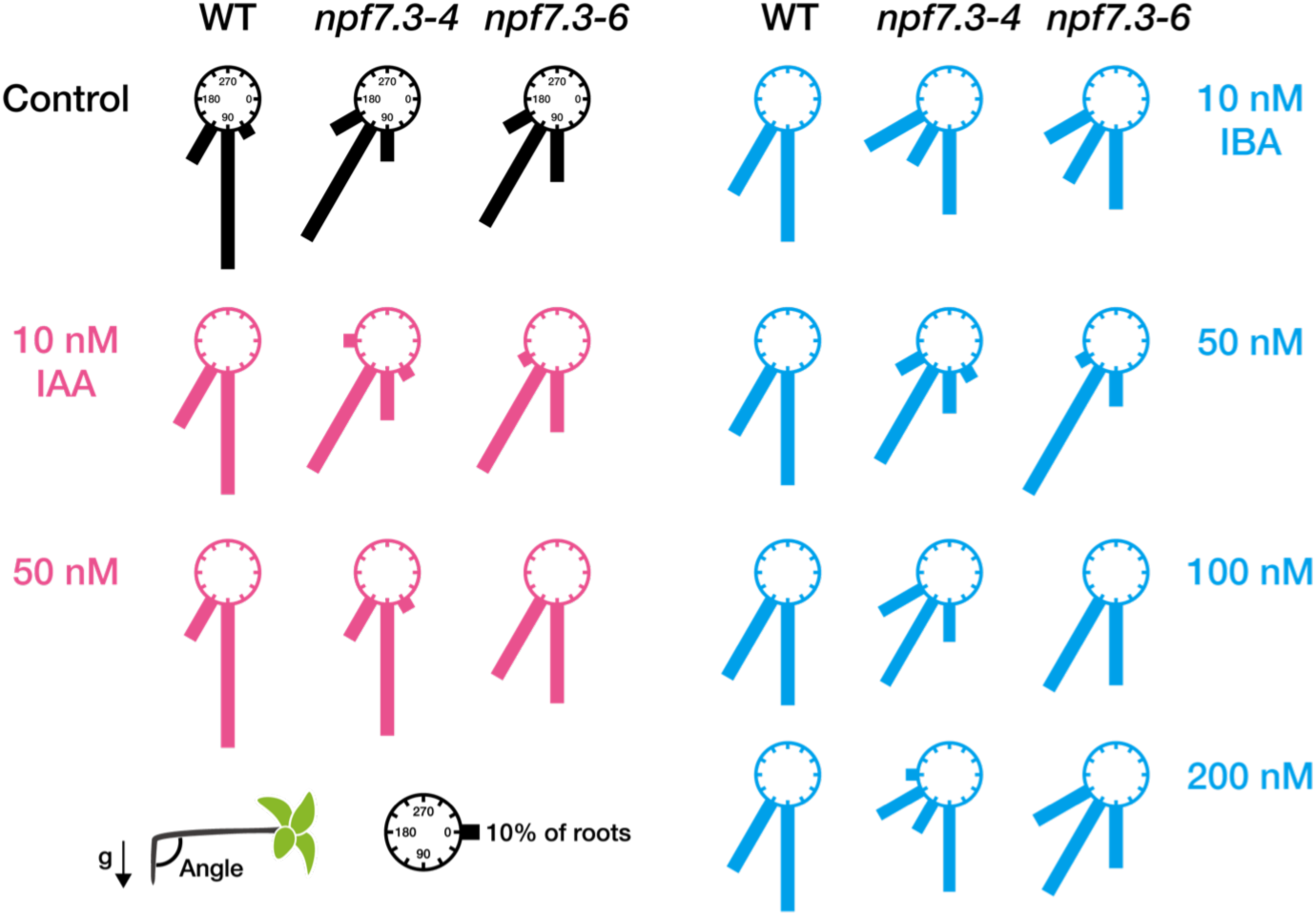
Effects of IBA and IAA on root gravitropism of n*pf7.3*. Wild type (WT) and *npf7.3* (*npf7.3-4* and *npf7.3-6*) seeds were germinated on control media or media containing IAA or IBA at the indicated concentrations. Plates were rotated 90° after seven days of cultivation at 22°C in continuous light. Primary root angles were measured one day after rotating the plates. Relative frequencies of root angles (30° intervals) are presented as the lengths of the bars (n = 10-20).

Although the bulk IAA content in root tissues was comparable between wild type and *npf7.3*, local auxin activities might be affected in *npf7.3* in the absence of cellular IBA uptake and, hence, subsequent IBA-to-IAA conversion. We first investigated the activities of an artificial auxin-inducible promoter *DR5* using a GFP or a GUS reporter (*DR5rev:GFP* and *DR5:GUS*, respectively). As reported previously, GFP fluorescence and GUS staining were observed in primary root tips and vascular tissues in the wild-type background grown on vertical plates (Fig. 5a, b). Similar expression patterns were observed in the *npf7.3* background, however, the signal intensities were significantly reduced in the mutant. We then examined how wild type and *npf7.3* respond to exogenous auxin in terms of *DR5* promoter activities (Fig. 5c). In the wild-type background, *DR5:GUS* expression was induced by both IBA and IAA in whole root tissues with higher expression levels in the root tip regions. *DR5:GUS* expression was induced by both IBA and IAA in the *npf7.3* background as well. However, the responses to IBA were much reduced in the mutant compared to wild type especially in root tips and elongation zones, whereas IAA treatment induced marker gene expression similarly in the wild-type and mutant backgrounds. We further analyzed the effects of root reorientation on the expression of *DR5rev:GFP*. When plants were grown on vertical plates, GFP signals were distributed symmetrically in the root tip regions of wild type and *npf7.3*, although lower signal intensities were observed for *npf7.3* (Fig. 5a). The distribution of GFP signals was changed in the wild-type background within four hours after rotating the plates 90° with higher intensities in the undersides, whereas an asymmetric distribution of GFP signals was not observed in the *npf7.3* mutant background (Fig. 5d). We also tested the effects of IBA and IAA on *DR5rev:GFP* expression (Fig. 5e). In the wild-type background, both IBA and IAA treatments enhanced auxin-responsive GFP signals and their asymmetric distribution in the root tip regions. IBA treatment, however, did not affect *DR5rev:GFP* expression, whereas IAA treatment induced asymmetric auxin responses in the *npf7.3* background. These results are consistent with the observation that IAA but not IBA treatment restored the disrupted root gravitropism observed in *npf7.3*.

**Fig. 5.**
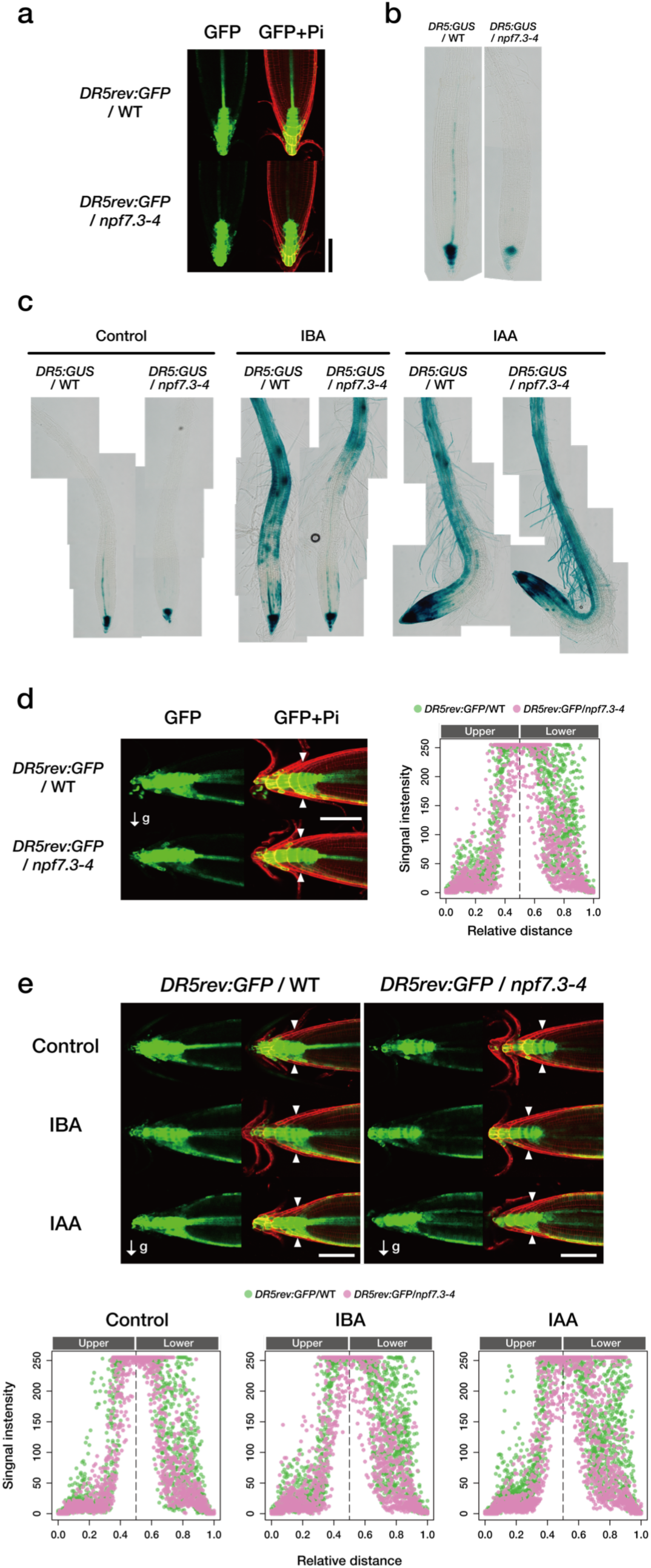
Auxin responses in *npf7.3*. **a**. Representative images of GFP fluorescence in root tips of seven-day-old wild-type and *npf7.3* expressing *DR5rev:GFP* (*DR5rev:GFP*/WT and *DR5rev:GFP*/*npf7.3-4*, respectively). Pi indicates propidium iodate staining of cell walls. Scale bar = 100 µm. **b,c**. Representative images of GUS staining patterns in the roots of seven-day-old wild-type and *npf7.3* expressing *DR5:GUS* (*DRrev5:GUS*/WT and *DR5rev:GUS*/*npf7.3-4*, respectively) before (**b**) and 24 hours after transferring to control media or media containing 4 µM IBA (IBA) or 200 nM IAA (IAA) (**c**). **d**. Representative images of GFP fluorescence 4 hours after rotating the plates (90°) containing seven-day-old wild-type and *npf7.3* expressing *DR5rev:GFP* (*DR5rev:GFP*/WT and *DR5rev:GFP*/*npf7.3-4*, respectively) (left panels). Pi indicates propidium iodate staining of cell walls. Scale bar = 100 µm. Intensities of GFP signals between the two arrowheads obtained from 19 and 20 plants, respectively, for *DR5rev:GFP*/WT and *DR5rev:GFP*/*npf7.3-4* are also shown (right panel). **e**. Representative images of GFP fluorescence 4 hours after rotating the plates (90°) containing seven-day-old wild type and *npf7.3* expressing *DR5rev:GFP* (*DR5rev:GFP*/WT and *DR5rev:GFP*/*npf7.3-4*, respectively) in the presence of IAA or IBA (upper panels). Plants were transferred to vertical plates containing IAA (200 nM) or IBA (4 µM) and then incubated for one hour before rotating the plates. Scale bar = 100 µm.

### Complementation of root gravitropic responses in *npf7.3* by NPF7.2 that has an IBA transport activity

NPF7.2, known as the low-affinity nitrate transporter NRT1.8, is the closest homologue of NPF7.3^28^. In many cases, homologous proteins have similar functions. Thus, we examined whether NPF7.2 could also transport IBA and found this to be the case (Fig. 6). NPF7.2 and NPF7.3 might play independent physiological roles in plants since their expression patterns are different (Supplementary Fig. 4)^21,29^. We reasoned that if IBA is the *in vivo* substrate of NPF7.3, NPF7.2 should be able to replace the function of NPF7.3 when expressed in the *npf7.3* mutant background under the control of the *NPF7.3* native promoter. Transgenic *npf7.3* plants expressing *NPF7.2* CDS under the control of a 2.1 kb promoter region of *NPF7.3* (*NPF7.3pro:NPF7.2*/*npf7.3*) showed gravitropic responses similar to wild type. In contrast, NPF4.6/NRT1.2/AIT1, an ABA transporter and also known as a low-affinity nitrate transporter, did not restore the impaired gravitropic responses observed in *npf7.3*.

**Fig. 6.**
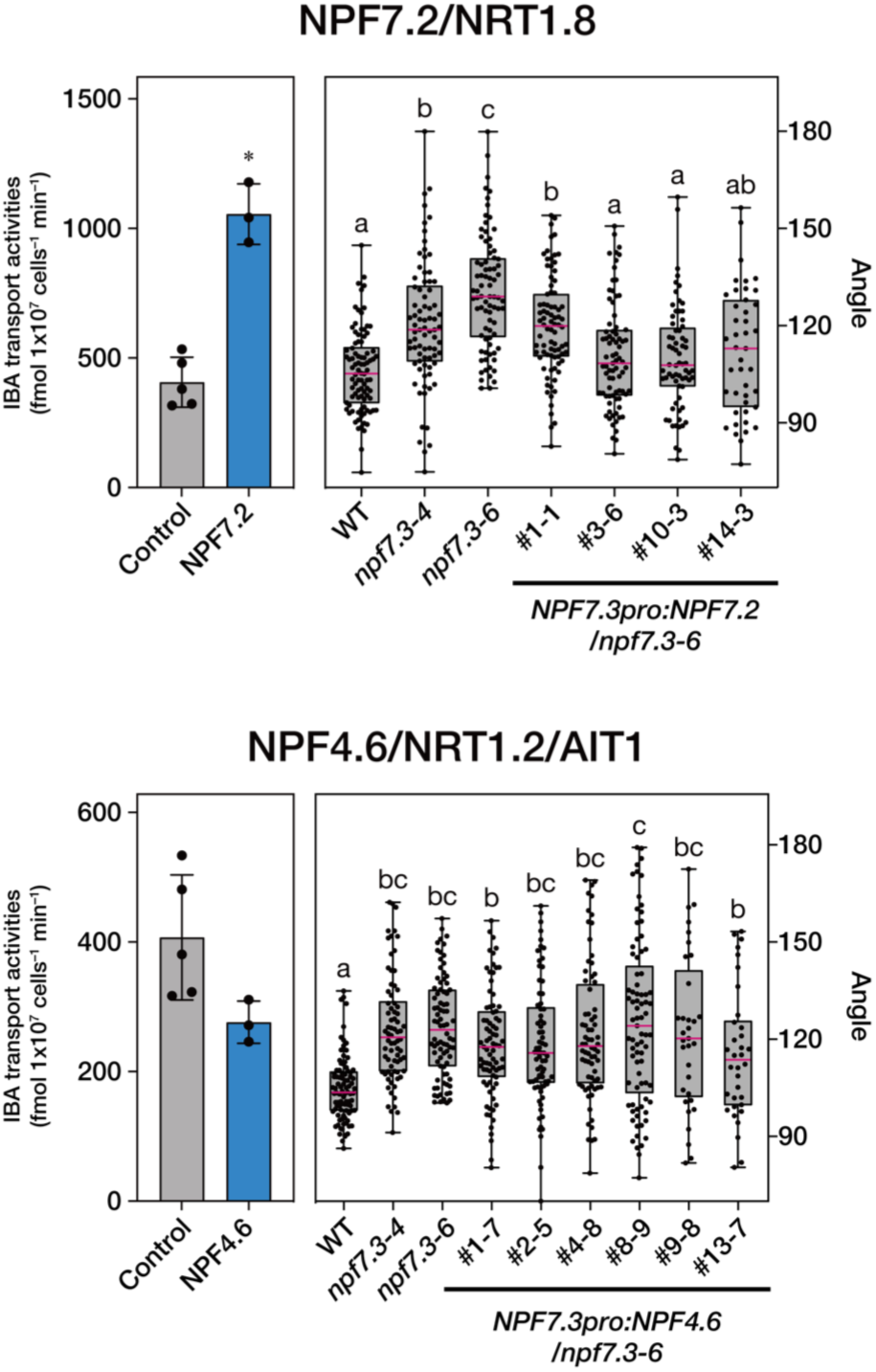
Complementation of *npf7.3* with another NPF protein having IBA transport activity. Activities of NPF7.2 and NPF4.6 to transport IBA were determined in yeast when the substrate concentration was 10 µM (left). Yeast cells containing the empty pYES2-DEST52 vector were analyzed as the control. Asterisks indicate significant differences compared with the controls (\**P* < 0.05 by Tukey’s multiple comparison test). Root angles of wild type (WT), *npf7.3* (*npf7.3-4* and *npf7.3-6*) and *npf7.3* expressing *NPF7.2* or *NPF4.6* under the control of the *NPF7.3* promoter (*NPF7.3pro:NPF7.2*/*npf7.3* and *NPF7.3pro:NPF4.6*/*npf7.3*, respectively) were measured 24 hours after the first rotation (right). Horizontal bars represent the median, boxes represent the middle 50% of the distribution, and whiskers represent the entire spread of the data. Different letters indicate statistically significant differences by Dunn’s multiple comparison test (*P* < 0.05).

### Expression patterns of *NPF7.3*

Previous studies have shown by in situ hybridization and histochemical analyses of promoter-GUS reporter lines that *NPF7.3* was expressed in the pericycle cells adjacent to xylem (xylem pole pericycle cells) in mature root tissues^19,30^. This observation is supported by high-resolution and genome-wide Arabidopsis gene expression profiling of root tissues (Supplementary Fig. 4a)^21,29^. The transcriptome analysis also suggested that *NPF7.3* is expressed in the stele of the root tip region as well. To investigate spatial expression patterns of *NPF7.3* in detail, we generated transgenic Arabidopsis expressing GUS or GFP fused to a nuclear localization signal (NLS) under the control of the *NPF7.3* promoter that was used for mutant complementation (Figs.1f and 6). In accordance with previous reports, strong GUS activities were detected in the xylem pole pericycle cells in mature root zones, however, we detected GUS staining also in the pericycle cells close to phloem tissues to a lesser extent (Fig. 7a, b). We further confirmed that GFP fluorescence was detected widely in pericycle cells (Fig. 7e, f). In addition, we detected GFP fluorescence and GUS activities in the stele of the root tip region and in columella root cap cells (Fig. 7c, d, g, h).

**Fig. 7.**
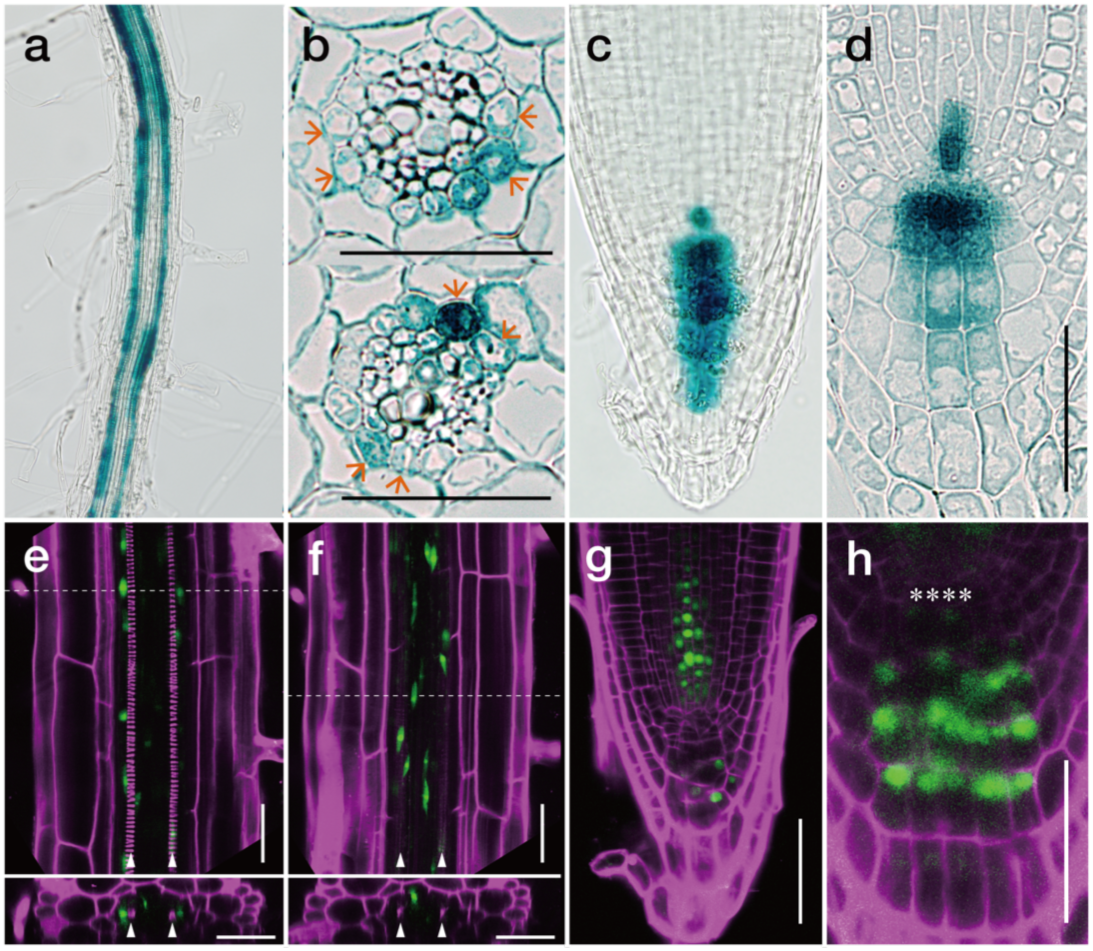
Spatial expression patterns of *NPF7.3* in roots. **a-d**. GUS-stained primary roots of *NPF7.3pro:GUS* transgenic plants. Seven-day-old seedlings grown on vertical plates were subjected to GUS staining at 37°C for 24 h. **a**. GUS activities in the mature root zone of a primary root. **b**. Transverse sections of the mature root zone of a primary root. Scale bars = 20 µm. Arrowheads indicate the positions of xylem pole pericycle cells. **c,d**. GUS activity in the root tip of a primary root. Scale bars = 50 µm. **e-h**. Representative images of GFP fluorescence in transgenic plants expressing GFP fused to a nuclear localization signal (NLD) under the control of NPF7.3 promoter (*NPF7.3pro:GFP-NLS*). **e,f**. Longitudinal sections (upper) and transverse sections (lower) of mature root tissues of the primary roots. Dashed lines in the upper panels indicate the positions of the transverse sections. **g,h**. Longitudinal sections of a primary root tip (**g**) and a magnified image of the root cap (**h**). T_1_ plants were selected on half-strength MS plates (placed horizontally) containing 25 mg/L hygromycin and 0.8% (w/v) agar for 5 days and transferred onto vertical plates without antibiotics. GFP fluorescence was observed after subsequent growth for seven days. GFP fluorescence and propidium iodate staining of cell walls are shown in green and magenta, respectively. Scale bars = 50 µm. Arrowheads indicate the positions of protoxylem. Asterisks indicate the position of quiescent center cells.

### Relationship with other IBA related mutants

The *ibr* mutants were isolated based on their reduced responsiveness to exogenous IBA; *ibr3* is defective in the IBA-to-IAA conversion whereas *ibr5* is impaired in auxin responses^31,32^. In contrast, *pen3* exhibits enhanced responses to IBA due to a defect in IBA extrusion from roots. We tested whether these mutants are also affected in their gravitropic responses, however, no significant differences in the root angles were observed compared to wild type (Supplementary Fig. 5). These results imply that IBA uptake into the pericycle cells and/or root tip cells mediated by NPF7.3 is the key process for generating local auxin concentration maxima to induce gravitropic responses and that other steps including the IBA-to-IAA conversion are rather redundant.

We also determined whether other IBA-mediated physiological responses are affected in *npf7.3* (Supplementary Fig. 6). That exogenous IBA treatment induces lateral root formation is well documented^33^. Although the number of lateral roots was comparable between wild type and *npf7.3* when grown on hormone-free media, *npf7.3* produced more lateral roots than wild type in response to IBA treatment. Furthermore, primary root elongation was less severely inhibited in *npf7.3* compared to wild type. Since the *npf7.3* response to IAA was similar to that of wild type, *npf7.3* is likely to be hypersensitive to IBA in lateral root formation as reported for *pen3*, but less sensitive to IBA in its inhibition of primary root elongation.

## Discussion

Although NPF proteins were originally identified as nitrate or di/tri-peptide transporters^15^, it is now evident that the protein family transports many different compounds. The presence of a relatively large number of family members in a given plant species (e.g. 53 homologues in Arabidopsis) indicates that their biochemical and physiological functions are diverse^16^. Upon cultivating several Arabidopsis mutants defective in NPFs, we first noticed a wavy root growth phenotype in *npf7.3* unlike that of wild type, (Fig. 1a-f). Wavy patterns of root growth are influenced by several environmental factors such as gravity. We found that *npf7.3* roots had a reduced ability to change their growth direction toward gravity compared to wild type (Fig. 1g, h). This observation is somewhat contradictory to previous studies since straight root growth with fewer wave numbers was reported to be associated with defective gravitropism and vice versa. Although the reason is unknown, it is possible that differences in growth conditions affected the phenotypes. In previous studies, plants were grown on inclined plates whereas we grew plants on vertical plates in this study. Also, we determined root gravitropic responses in the absence of sucrose, a media component normally present in previous studies. In any case, our observations indicate that NPF7.3 plays an important role in root gravitropism. NPF7.3 has been characterized previously as a nitrate and potassium transporter^19,20^. However, it is not likely that the altered root gravitropism observed in *npf7.3* was caused by defects in nitrate and/or potassium transport since the phenotype was seen even when the nutrients were abundant in the media. Instead, we hypothesize that NPF7.3 somehow regulates auxin responses, since the hormone plays a critical role in gravitropism.

IAA is the major naturally occurring auxin. Recent studies have demonstrated that the majority of IAA present in plants is synthesized from tryptophan via reactions catalyzed by TAA/TAR and YUCCA proteins^34^. IBA was previously recognized as a minor endogenous auxin since its presence was reported in several plant species, and application of IBA to plants induced “auxin-like” responses. More recent studies, especially characterization of mutants with altered responses to exogenous IBA, revealed that IBA must be converted to IAA via peroxisomal ß-oxidation to induce physiological responses^31,35,36^. This theory is also supported by the fact that IBA is not bound to the auxin receptor TIR1/ABFs^37,38^. Thus, IBA is now considered to be a precursor of IAA, although how IBA is synthesized in plants is unknown. Interestingly, one of the Arabidopsis NPF proteins, NPF6.3, transports IAA *in vivo* besides its well-known function as the nitrate transporter NRT1.1/CHL1^17,39^. More recently, another NPF protein, NPF5.12 was shown to be an IBA transporter (also called TOB1)^40^. Therefore, it is possible that NPF7.3 transports IAA, IBA and/or their related compounds. Our biochemical analyses using yeast as a heterologous expression system showed that NPF7.3 mediates the cellular uptake of IAA and IBA (Fig. 2). Consistent with this result, a previous study reported that NPF7.3 was localized to the plasma membrane in plant cells^19^. In contrast, NPF5.12/TOB1 was localized to vacuoles and regulates the IBA content in subcellular compartments. Some NPF proteins, including NPF7.3/NRT1.5, mediate nitrate export from inside cells to the outside. If an IAA or IBA exporter were expressed in our yeast system, the cells should accumulate a smaller amount of the compound compared to the control cells; however, this result was not the case for NPF7.3. NPFs are generally proton symporters. In accordance with this detail, IAA and IBA uptake activities were relatively high when the buffer was acidic (Fig. 2a, b), although we cannot exclude the possibility that NPF7.3 recognizes protonated IAA and IBA as substrates. The pH values for the apoplast and cytosol are estimated to be 5.0 to 6.0 and 7.2 to 7.4, respectively^41,42^. Therefore, it is reasonable to propose again that NPF7.3 is an IAA/IBA importer. IAA and IBA are weak acids (p*K*a = 4.75 and 4.83, respectively) with a carboxylic group and exist as protonated or ionized forms depending on pH^43^. Their protonated forms can passively move through biological membranes relatively easily because of their hydrophobic natures. Thus, one may imagine that specific transporters are not required especially when IAA and IBA are taken into cells; however, at least for IAA uptake, the existence of specific transporters is evident. Fig. 2 shows that IBA is less permeable to biological membranes compared to IAA. This observation suggests that cellular IBA uptake is also mediated by specific transporters. In our yeast transport assays, NPF7.3 transported IBA more efficiently than IAA, although the *K*m values for the two substrates were comparable (Fig. 2c). Thus, it is possible that IBA is the preferred substrate of NPF7.3 *in vivo*, even though endogenous IBA levels are significantly lower than IAA levels (Fig. 3).

As mentioned above, IBA is converted to IAA^31,35,36^. We therefore hypothesized that IBA taken into cells mediated by NPF7.3 is converted to IAA to induce root gravitropic responses. If this hypothesis is correct, exogenous application of IAA would complement the phenotype observed in *npf7.3*. In contrast, IBA treatment would be less effective in rescuing the *npf7.3* phenotype since the mutant is impaired in IBA uptake into cells and, thus, not able to convert IBA to IAA efficiently. As expected, IAA, but not IBA, was able to restore the altered gravitropic responses in the *npf7.3* mutant roots (Fig. 4). These data suggest that auxin responses are reduced in *npf7.3*. Although bulk IAA content in whole root tissues was comparable between wild type and *npf7.3* (Fig. 3), it is still possible that IAA levels are affected very locally. We observed that auxin responses determined by *DR5rev:GFP* and *DR5:GUS* reporters were significantly lower in the root tip region of *npf7.3* compared to wild type (Fig. 5a, b). Although *DR5:GUS* expression in root tissues was similarly induced by exogenous IAA in wild-type and *npf7.3* backgrounds, the reporter had a lower response to IBA in *npf7.3* than the wild type (Fig. 5c). In addition, we found that the asymmetric distribution of auxin activities observed after reorientation of the roots was not created in the *npf7.3* mutant background (Fig. 5d). Furthermore, *npf7.3* roots treated with IAA, but not with IBA, showed asymmetric expression patterns of *DR5rev:GFP* upon reorientation (Fig. 5e). Finally, we confirmed that the expression of an NPF protein having an IBA transport activity (NPF7.2) in the *npf7.3* background under the control of the *NPF7.3* promoter complemented the *npf7.3* phenotype (Fig. 6). Collectively, these data consistently support the idea that NPF7.3 mediates cellular IBA uptake in root tissues, and a defect in this process results in a reduced ability to synthesize IAA from IBA and to establish proper auxin (IAA) gradients within roots in response to gravity.

Many textbooks illustrate that IAA is actively synthesized in the shoot apex and transported toward the root tips via phloem tissues and/or cell-to-cell polar transport mechanisms. Recent studies, however, have demonstrated that IAA synthesized locally in roots also plays important roles in root growth and development^44,45^, even though it is largely unknown how IBA is transported within plants. Given that IBA levels are reduced in *npf7.3* roots compared to wild type, NPF7.3 might play a role in retaining IBA in root tissues and/or IBA transport from shoots to roots. In this study, we found that *NPF7.3* was expressed predominantly in pericycle cells in mature root tissues whereas we were unable to detect its presence in above ground tissues (Fig. 7a, b, e, f). Perhaps IBA is normally transported from roots to shoots via xylem in wild type, and NPF7.3 facilitates IBA uptake into pericycle cells from xylem (Fig. 8a). In fact, IBA was reported to be transported from roots to shoots at least when applied exogenously^46^. In this case, the loss of NPF7.3 function would increase the amount of IBA translocated from roots to shoots and, thus, bulk IBA levels in the roots would be reduced. This hypothesis is not fully proven yet since we were unable to detect IBA in shoot tissues possibly due to impurities affecting liquid chromatography tandem-mass spectrometry (LC-MS/MS) analysis and/or rapid degradation of IBA in shoots. Interestingly, we also found that *NPF7.3* was expressed in the root tip region (stele cells and columella root cap cells) of primary roots (Fig. 7c, d, g, h). Columella root cap cells are considered to be important hubs to perceive and transmit gravity signals and to generate asymmetric IAA distribution^11^. IBA-to-IAA conversion is active in the same region^32,47^, indicating that NPF7.3 is required for efficient IAA production from IBA. IAA synthesized from IBA could be exported from columella root cap cells upon reorientation, thereby rapidly inducing an auxin response on the gravity side (Fig. 8b). Again, the site of IBA synthesis is not known. However, we propose that part of the IBA taken into pericycle cells from xylem is transported to the root tip regions through phloem tissues and/or other unknown transport mechanisms. It is also possible that IBA is converted to IAA in pericycle cells and then transported to the root tip through phloem tissues and/or polar transport systems. Previous studies have also indicated that IBA is exported from lateral root cap cells toward the rhizosphere mediated by the ABC type transporters ABCG36/PEN3 and ABCG37/PIS1^13^. Therefore, if NPF7.3 is not present in the root tip region, more IBA would be extruded from the roots (Fig. 8b). This possibility could also explain the reduced IBA levels in *npf7.3* roots.

**Fig. 8.**
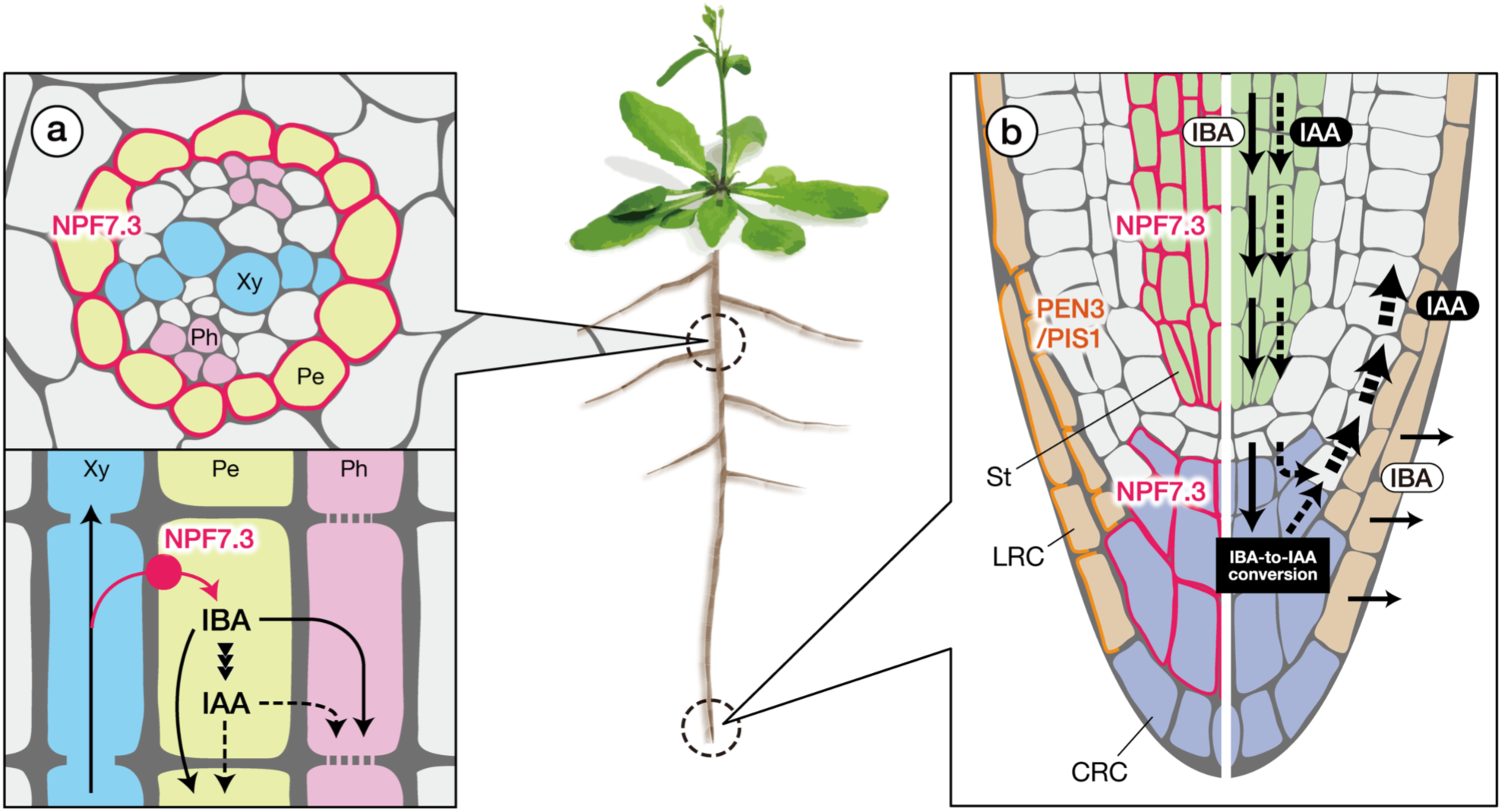
Possible physiological functions of NPF7.3 in roots. In mature root tissues, NPF7.3 is produced in pericycle cells. Given that IBA levels are reduced in *npf7.3* roots, it is possible that NPF7.3 is involved in the uptake of IBA from xylem (**a**). IBA and/or IAA converted from IBA is then transported toward the root tip region through phloem tissues and/or basipetal transport mechanisms. *NPF7.3* expression in the root tip region suggests that cellular IBA uptake is also active in this region (**b**). IBA-to-IAA conversion is proposed to occur in the root tip region, especially in root cap cells. IAA synthesized from IBA is unevenly distributed within the root tip via polar transport mechanisms to induce asymmetric cell elongation in response to gravity. In *npf7.3*, the ability of root cap cells to retain IBA is reduced, thereby resulting in lower IBA levels in the roots due to continuous IBA extrusion mediated by PEN3 and PIS1 (also referred to as ABCG36 and ABCG37, respectively). Basipetal transport of IBA and/or IAA is also reduced in *npf7.3* as explained in (**a**). Pink and orange lines represent the localization of NPF7.3 and PEN3/PIS1, respectively. Solid and dashed arrows indicate the predicted flows of IBA and IAA, respectively. The stele is indicated in green. CRC, columella root cap (indicated in purple); LRC, lateral root cap (indicated in orange); Pe, pericycle; Ph, phloem; St, stele; Xy, xylem.

In contrast to *npf7.3*, mutants defective in IBA-to-IAA conversion (*ibr3-9*), had a response to gravity similar to wild type (Supplementary Fig. 5), probably because the IBA-to-IAA conversion mediated by IBR3 is redundant. In fact, mutants defective in several acyl-CoA oxidases are insensitive to IBA^48^. Likewise, enzymes involved in other steps of the IBA-to-IAA conversion also have homologues^35^. Never the less, it is likely that cellular IBA uptake mediated by NPF7.3 is specific. We showed that NPF7.2 also has an IBA transport activity, however, the gene is not expressed in the root tip region according to a public database (Supplementary Fig. 4)^21^. Thus, we think that cellular IBA uptake in the root tip region is the key process for regulating the local distribution of IAA and, hence, root gravitropic responses. In addition to the defect in gravitropism, *npf7.3* showed altered responses to exogenous IBA treatment. Consistent with the observation that *npf7.3* is less sensitive to IBA (Fig. 4), primary root elongation of *np7.3* was less inhibited by IBA than wild type (Supplementary Fig. 6b). Nevertheless, *npf7.3* produced a larger number of lateral roots than wild type in response to IBA treatment. IBA synthesized in the lateral root cap cells rather than in the columella cells is proposed to be important for inducing lateral root formation. Since *NPF7.3* is expressed in columella cells, possibly the loss of NPF7.3 function resulted in increased IBA-to-IAA conversion in the lateral root cap cells and, hence, enhanced lateral root formation.

To our knowledge, our study is the first demonstration of the role of endogenous IBA in regulating root gravitropic responses. Why the IBA pathway is required to fully induce gravitropism in addition to the main IAA biosynthesis pathway remains an enigma. Furthermore, it will be important to establish how IBA is synthesized in plants. A public database indicates that the expression of *NPF7.3* is regulated by biotic and abiotic stresses. Thus, it will also be informative to know whether the expression of *NPF7.3* is related to IBA transport. If not, what is the substrate of NPF7.3 and what is the physiological role of the protein under stressed conditions? These questions will be answered in future studies.

## Methods

### Plant materials and growth conditions

Arabidopsis [*Arabidopsis thaliana* (L.) Heynh.] accession Columbia-0 (Col-0) was used as the wild-type for all experiments in this work. Mutant lines obtained from the Arabidopsis Biological Resource Center used in experiments were as follows: *npf7.3-4* (*nrt1.5-4*) (SALK_063393; Li *et al*.)^28^, *npf7.3-6* (GK-877E12, newly identified in this study; see Supplementary Fig. 1), *ibr3-9* (SALK_033467)^31^, *ibr5-3* (SALK_039359)^49^ and *pen3-4* (SALK_000578)^50^. A homozygous *npf7.3-6* mutant was selected by PCR using primer combinations designed by the T-DNA Primer Design Tool (Table S1; http://signal.salk.edu/tdnaprimers.2.html). Transgenic *npf7.3* lines expressing *DR5:GUS* or *DR5rev:GFP* were generated by crossing *npf7.3-4* with *DR5:GUS*/WT^51^ or *DR5rev:GFP*/WT^52^, respectively. *DR5:GUS*/WT was a gift from Dr. Tom J. Guilfoyle, and *DR5rev:GFP*/WT was obtained from the Nottingham Arabidopsis Stock Centre. After surface sterilization with 70% (v/v) ethanol and then with 5% (v/v) NaClO (0.25% active chlorine) containing 1% (w/v) SDS, seeds were sown on half-strength MS media containing 1.5% (w/v) agar. Sucrose was not added to the media unless otherwise indicated. To break dormancy, seeds were incubated at 4°C for 3 days in the dark, and the plates were vertically placed in growth chambers at 22°C under continuous light (50-70 µmol photons m–2 s–1).

### Transport assay in yeast cells

IBA and IAA import activities were determined by analyzing the compounds taken into yeast cells expressing NPF proteins using LC-MS/MS as described previously^53^. Previously constructed NPF CDSs^54^ in the yeast expression vector pYES-DEST52 (Thermo Fisher) were introduced into the *Saccharomyces cerevisiae* strain INVSc1 (Thermo Fisher). The empty pYES-DEST52 vector was transformed as a negative control. Yeast cells containing the pYES-DEST52 constructs were precultured in SD media lacking uracil (SD, -Ura) overnight. The precultured cells were collected by centrifugation and then resuspend with SG media containing 2% (w/v) galactose and 1% (w/v) raffinose without uracil (SG, -Ura) to induce the production of NPF proteins. After centrifugation, yeast cells were resuspended in 50 mM potassium phosphate buffer (KPB) (pH 5.8 or 7.0) and incubated with the buffer containing IBA or IAA at 25°C for 10 min. For competition assays, the cells were incubated with IBA and either potassium nitrate or IAA. Concentrations of the substrates are indicated in the figure legends. Yeast cells were then collected by centrifugation and followed by washing three times with 50 mM KPB (pH 7.0). We defined the end of the reactions as the first wash with KPB (pH 7.0). The cells were stored at –80°C until extraction. Quantification of IBA and IAA by LC-MS/MS was performed as described below.

### Hormone measurements

To extract IBA and IAA from plant tissues, lyophilized samples were homogenized by vortexing with zirconia beads in 80% (v/v) acetonitrile containing 1% (v/v) acetic acid and isotope-labeled IBA ([^13^C_8_, 15N_1_]IBA) and/or IAA ([methylene-^2^H_2_]IAA) as internal standards. Homogenates were incubated in the darkness for 16 h at 4 °C and then centrifuged at 3,000 × *g* for 5 min at 4 °C. The supernatants were dried under N_2_ flow at 40 °C. The samples were resuspended in 1% (v/v) acetic acid by vortexing and sonication and then applied onto Oasis HLB cartridges (Waters) that were conditioned with acetonitrile and 1% (v/v) acetic acid. After washing with 20% acetonitrile containing 1% (v/v) acetic acid, fractions containing IBA and IAA were eluted with 60% acetonitrile containing 1% (v/v) acetic acid. The eluates were dried under N_2_ flow, and the precipitates were resuspended with 1% (v/v) acetic acid. The suspensions were loaded onto Oasis WAX cartridges (Waters) pre-conditioned with acetonitrile, 0.1M potassium hydrate and 1% (v/v) acetic acid, followed by washing with 20% (v/v) acetonitrile containing 1% (v/v) acetic acid. IBA and IAA were eluted with 60% (v/v) acetonitrile containing 1% (v/v) acetic acid and then completely dried under N_2_ flow. The extracts were dissolved with 50 µl 1% (v/v) acetic acid and subjected to LC-MS/MS analysis.

To extract IBA and IAA from yeast cells, frozen samples were incubated for 2 h at 4°C in 80% (v/v) acetone containing 1% (v/v) acetic acid and [methylene-^2^H_2_]IBA and/or [methylene-2H_2_]IAA as internal standards. Supernatants were collected by centrifugation at 20,000 × *g* for 5 min and then completely dried under N_2_ flow at 40°C. The samples were resuspended in 1% (v/v) acetic acid by vortexing and sonication and then loaded onto Oasis WAX cartridges that were conditioned with 1% (v/v) acetic acid, followed by washing with acetonitrile containing 1% (v/v) acetic acid. IBA and IAA were eluted with 80% (v/v) acetonitrile containing 1% (v/v) acetic acid and then completely dried under N_2_ flow. The extracts were dissolved in 50 µl 1% (v/v) acetic acid and subjected to LC-MS/MS analysis.

In the experiments presented in Supplementary Fig. 3, IAA, IAA-amino acid conjugates [IAA-aspartate (IAA-Asp) and IAA-glutamate (IAA-Glu)], and 2-oxoindole-3-acetic acid (oxIAA) were extracted and purified as described previously^55^. In this analysis, [phenyl-^13^C_6_]IAA was used as an internal standard instead of [methylene-^2^H_2_]IAA.

Endogenous ABA, jasmonic acid (JA), JA-Ile, *trans*-zeatin (tZ), isopentenyl adenine (iP) and salicylic acid (SA) were extracted and purified as described previously^53^.

IAA-Asp, IAA-Glu and oxIAA were analyzed using an Agilent 6420 Triple Quad LC-MS/MS system (Agilent) equipped with a ZORBAX Eclipse XDB-C18 column. Conditions for LC and MS/MS analyses were the same as described previously^55^. For analyzing other chemicals, a Nexera HPLC system (Shimadzu) coupled with a quadrupole/time-of-flight tandem mass spectrometer (Triple TOF 5600, AB Sciex) was used. Conditions for LC and for MS/MS analyses of IBA and IAA are summarized in Supplementary Tables 2 and 3. LC and MS/MS conditions for other chemicals were the same as previously described^53^.

### Chemicals

IBA, IAA, [methylene-^2^H_2_]IAA and ABA were purchased from Sigma-Aldrich. [^13^C_8_,^15^N_1_]-IBA was synthesized according to the literature^56^ from [^13^C_8_,^15^N_1_]-indole (Cambridge Isotope Laboratories, CNLM-4786-0.1) and was purified by silica gel column chromatography (CHCl_3_-MeOH=10:1). The purity of isolated [^13^C_8_,^15^N_1_]-IBA was determined by HPLC in comparison with authentic IBA. [methylene-^2^H_2_]IBA was purchased from CDN Isotopes. [phenyl-^13^C_6_]oxIAA, [^13^C_4_,^15^N]IAA-Asp and [^13^C_5_,^15^N]IAA-Glu were previously synthesized^34,57^. JA, JA-Ile and [^13^C_6_]JA-Ile^58^ were gifts from Dr. Yusuke Jikumaru (RIKEN). [^2^H_2_]JA was purchased from Tokyo Chemical Industry Co. [^2^H_2_]ABA was purchased from Icon Isotope. [^2^H_5_]tZ and [^2^H_6_]iP were purchased from OlChemIm. [^2^H_6_]SA was purchased from Isotec.

### Vector construction and generation of transgenic plants

Sequences of primers used for vector construction in this study are listed in Supplementary Table 1. For complementation of *npf7.3*, transgenic *npf7.3* containing either *NPF7.3pro:NPF7.3, NPF7.3pro:NPF7.2*, or *NPF7.3pro:NPF4.6* constructs were generated. A 2.1 kb sequence containing the *NPF7.3* promoter was amplified from Arabidopsis genomic DNA by PCR using primers NPF7.3pro-topo-F and NPF7.3pro-R and cloned into pENTR/D-TOPO (pENTR-*NPF7.3pro*) (Invitrogen). The *NPF7.3* promoter sequence was amplified again by PCR from the pENTR-*NPF7.3pro* construct using primer pairs NPF7.3pro-attL1-IF-F/NPF7.3pro-NPF7.3CDS-IF-R, NPF7.3pro-attL1-IF-F/NPF7.3pro-NPF7.2CDS-IF-R, and NPF7.3pro-attL1-IF-F/NPF7.3pro-NPF4.6CDS-IF-R and then cloned into the pDONR207 vector containing *NPF7.3*, the pENTR vector containing NPF7.2 or the pENTR vector containing NPF4.6^54^, respectively, using an In-Fusion Cloning Kit (Takara Bio). The vectors were linearized by inverse PCR using primer pairs NPF7.3CDS-IF-F/pDONR-attL1-R (pDONR207-*NPF7.3*), NPF7.2CDS-IF-F/pENTR-attL1-R (pENTR-NPF7.2) and NPF4.6CDS-IF-F/pENTR-attL1-R (pENTR-*NPF4.6*). The *NPF7.3pro:NPF* sequences were introduced into the pGWB1 vector^59^ by LR reactions. The final binary vectors were introduced into *Agrobacterium tumefaciens* strain GV3101 by electroporation and used to transformed the *npf7.3-6* mutant by the floral dip method as previously described^60^.

To generate the *NPF7.3pro:GUS* line, the 2.1 kb *NPF7.3* promoter sequence cloned in pENTR-*NPF7.3pro* was amplified by PCR using primers NPF7.3pro-IF-F and NPF7.3pro-GUS-IF-R. The *GUS* gene was amplified from pGWB3 by PCR using primers GUS-F and GUS-IF-R. The *NPF7.3pro* and *GUS* fragments were combined and then cloned using an In-Fusion Cloning Kit into the linear pENTR vector amplified from pENTR/D-TOPO by inverse PCR using primers pENTR-F and pENTR-R. The *NPF7.3pro:GUS* sequence was then cloned into the pGWB501 vector^59^ by the LR reaction. The final binary vector was introduced into GV3101 by electroporation and transformed into plants by the floral dip method.

To generate transgenic plants expressing GFP fused to SV40 NLS, the *NLS* sequence was generated by annealing complementary pairs of oligonucleotides, 5’-TCCGGACTCAGATCTCGAGCTGATCCAAAAAAGAAG-3’/5’-CTTTCTCTTCTTTTTTGGATCAGCTCGAGATCTGAGTCCGGA-3’, 5’-TCCGGACTCAGATCTCGAGCTGATCCAAAAAAGAAG-3’,5’-CTTTCTCTTCTTTTTTGGATCAGCTCGAGATCTGAGTCCGGA-3’, and 5’-TCCGGACTCAGATCTCGAGCTGATCCAAAAAAGAAG-3’/5’-CTTTCTCTTCTTTTTTGGATCAGCTCGAGATCTGAGTCCGGA-3’. The annealed fragments were ligated by T4 DNA ligase to create a tandemly repeated *NLS* sequence. The *NLS* sequence was further amplified from the ligation mixture by PCR using primers GFP-NLS-IF-F and GFP-NLS-IF-R. The *GFP* gene was amplified from the pGWB5 vector^59^ by PCR using primers GFP-IF-F and GFP-IF-R. The amplified *GFP* and *NLS* fragments were cloned using an In-Fusion Cloning Kit into pENTR4 digested with NcoI and XhoI to create pENTR4-GFP-NLS. The *GFP*-*NLS* sequence was amplified from pENTR4-GFP-NLS by PCR using primers NRT7.3pro-GFP-NLS-IF-F and NRT7.3pro-GFP-NLS-IF-R. The PCR fragments were cloned using an In-Fusion Cloning Kit into the linear pENTR-NPF7.3pro vector amplified by inverse PCR using primers pENTR-NPF7.3pro-F and pENTR-NPF7.3pro-R. The construct was cloned into the pGWB501 vector^59^ by LR reactions. The final recombinant binary vector was introduced into GV3101 by electroporation and transformed into plants by the floral dip method.

### Histochemical GUS analysis

GUS activity was detected using a staining solution composed of 50 mM sodium phosphate buffer (pH 7.2), 10 mM EDTA, 0.05% (v/v) Triton X-100, 0.5 mM potassium ferrocyanide, 0.5 mM potassium ferricyanide and 1 mM X-Gluc. Samples were mounted in chloral hydrate after stopping the reaction with 70% (v/v) ethanol and then observed using an Olympus BX53 microscope (Olympus).

For the preparation of root sections, GUS-stained samples were fixed with 0.05 M sodium cacodylate buffer (pH 7.4) containing 4% (w/v) paraformaldehyde and 2% (v/v) glutaraldehyde overnight at 4°C. Fixed root samples were dehydrated through a graded ethanol series and embedded in Technovit® 7100 resin (Kulzer). Thin sections (1 µm thick) were obtained using a glass knife on an ultramicrotome (EM UC7; Leica) and then observed using an Olympus BX53 microscope.

### Microscopic observations of GFP signals

GFP-fluorescent images were obtained using a confocal laser scanning microscope LSM700 (Carl Zeiss). Root tips were excised from whole seedlings and then stained with a 30 µM µµdium iodide (Pi) solution to visualize cell walls. Spatial distributions and intensities of GFP signals were quantified using the ZEN software (Carl Zeiss).

### Statistical analysis

Significant differences between two groups were determined by Student’s *t*-test. Significant differences among three or more groups were evaluated by Tukey’s or Dunn’s multiple comparison test. All statistical tests were performed using statistical software Prism 8 (GraphPad Software Inc., California, USA) or R ver. 3.6.1 (R Core Team) in RStudio ver. 1.2.5001 (RStudio Team).

## Acknowledgements

We thank Dr. Tsuyoshi Nakagawa (Shimane University) for providing the pGWB1, pGWB5 and pGWB501 vectors; and Dr. Tom J. Guilfoyle (University of Missouri) for providing the *DR5:GUS*/WT seeds. We also thank Ms. Tomoe Nose and Masako Tanaka (RIKEN) for their technical assistance. This study was supported in part by a Grant-in-Aid for Scientific Research (no. 17H06470 and 17H06477 to M.U. and no. 19K06708 to N.T.) from the Japan Society for the Promotion of Science (JSPS) and the RIKEN Special Postdoctoral Researcher Program to S.W.

## Author contributions

S. W. and M.S. designed the study. S.W. performed genotyping and phenotypic analyses of mutants. S.W. and Y.K. cloned and constructed vectors for transport assays and mutant complementation. Y.K. conducted transport assays. Y.K., Y.A. and H.K. analyzed plant hormones. K.H. synthesized the isotope-labeled IBA. S.W., H.S., N.T. and M.U. generated and observed transgenic lines expressing *DR5rev:GFP* and *DR5:GUS*. S.W., N.T-K. and K.T. performed GUS staining and prepared cross sections. S.W., N.T., H.K., K.H., M.U. and M.S. wrote the manuscript. All authors discussed the results and commented on the manuscript.

## Competing interests

The authors declare no competing interests.

## Supplementary information

Supplementary Figs. 1-6, Tables 1-3, and References

## Notes

### Competing Interest Statement

The authors have declared no competing interest.

