## Supplementary Information for "The Arabidopsis NRT1/PTR FAMILY Protein NPF7.3/NRT1.5 is an Indole-3-butyric Acid Transporter Involved in Root Gravitropism"

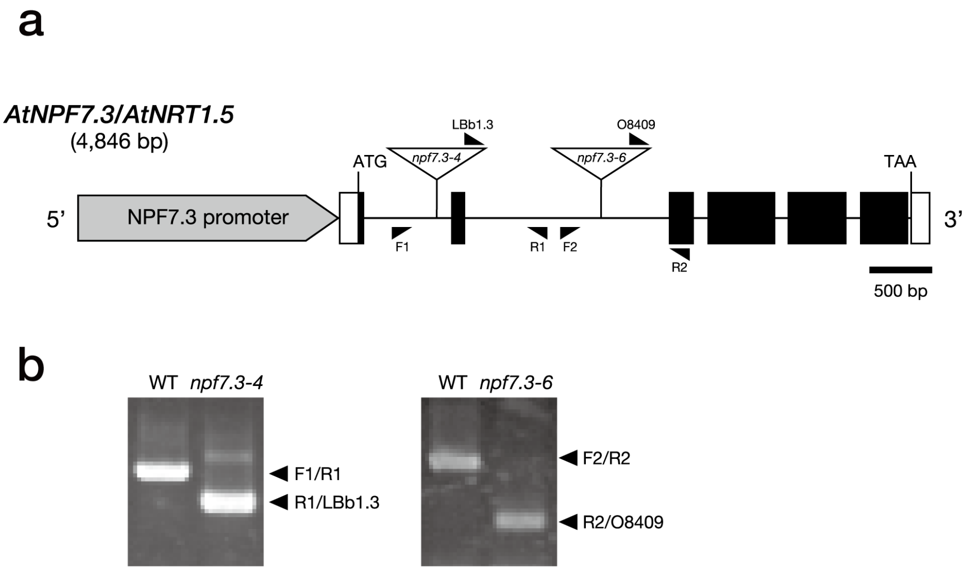


**Supplementary Fig. 1. T-DNA insertion mutants of *NPF7.3/NRT1.5* used in this study. a**. Structure of the *NPF7.3/NRT1.5* gene and positions of T-DNA insertions in two *npf7.3* alleles. Arrowheads indicate the positions of primers used for PCR-based genotyping of *npf7.3-4* (SALK_063393) and *npf7.3-6* (GK-877E12). **b**. Confirmation of homozygous T-DNA insertions by PCR.

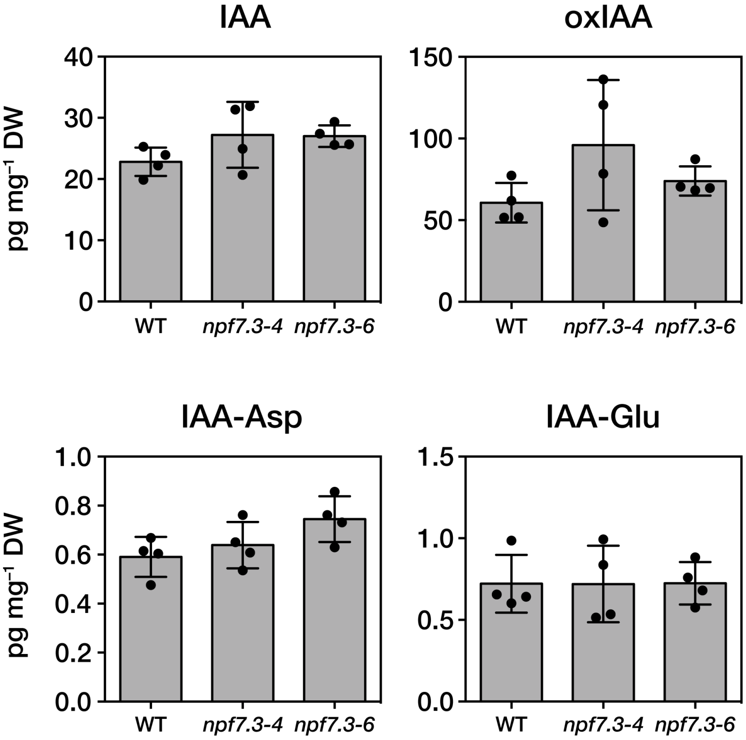


**Supplementary Fig. 2. Endogenous levels of IAA derivatives in *npf7.3*.** IAA, 2-oxoindole-3-acetic acid (oxIAA), IAA-aspartate (IAA-Asp) and IAA-glutamate (IAA-Glu) were quantified from the roots of 14-day-old wild type (WT) and *npf7.3* (*npf7.3-4* and *npf7.3-6*) seedlings. Bars indicate the standard deviations of the means. Dots represent individual measurements. Asterisks indicate significant differences compared with wild-type (**P* < 0.05 by Tukey's multiple comparison test).

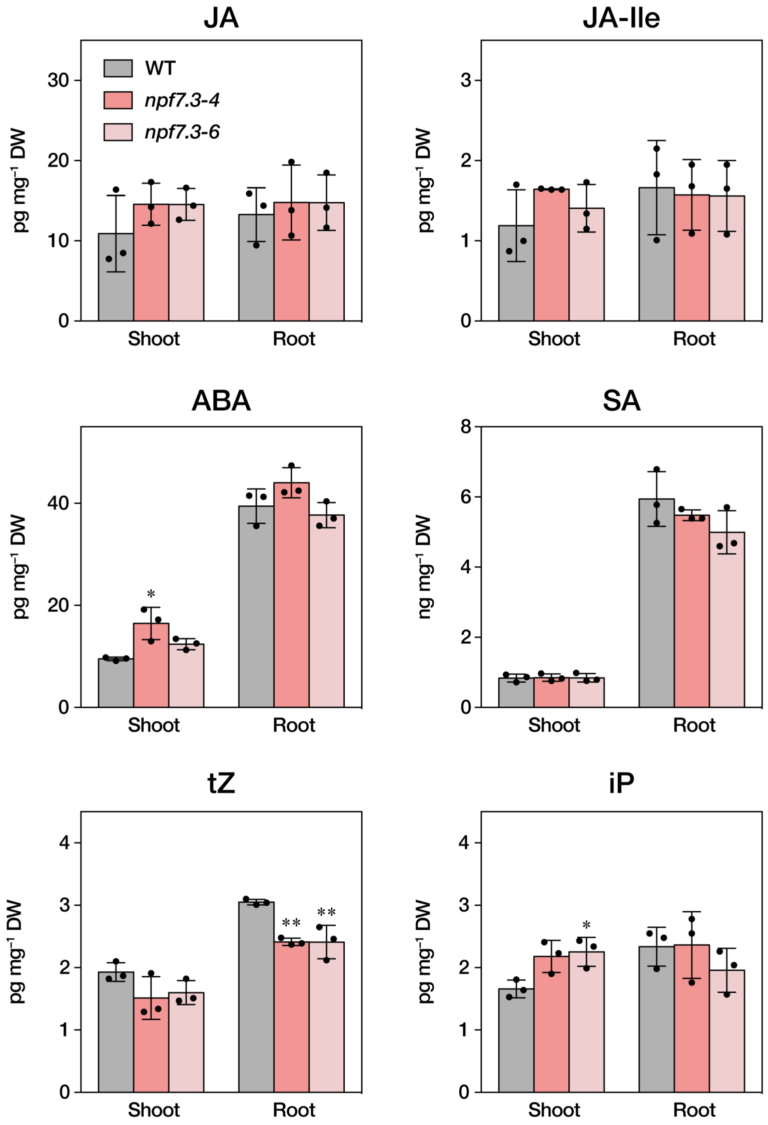


**Supplementary Fig. 3. Endogenous plant hormone levels in *npf7.3*.** Endogenous levels of jasmonic acid (JA), JA-Ile, ABA, salicylic acid (SA), *trans*-zeatin (tZ) and isopentenyl adenine (iP) were measured in shoot and root tissues excised from 14-day-old wild-type and *npf7.3* (*npf7.3-4* and *npf7.3-6*). Bars indicate standard deviations of the means. N.D, not detected. Asterisks indicate significant differences as compared with wild type within each treatment group (*$P$ < 0.05; **$P$ < 0.01 by Tukey's multiple comparison test).

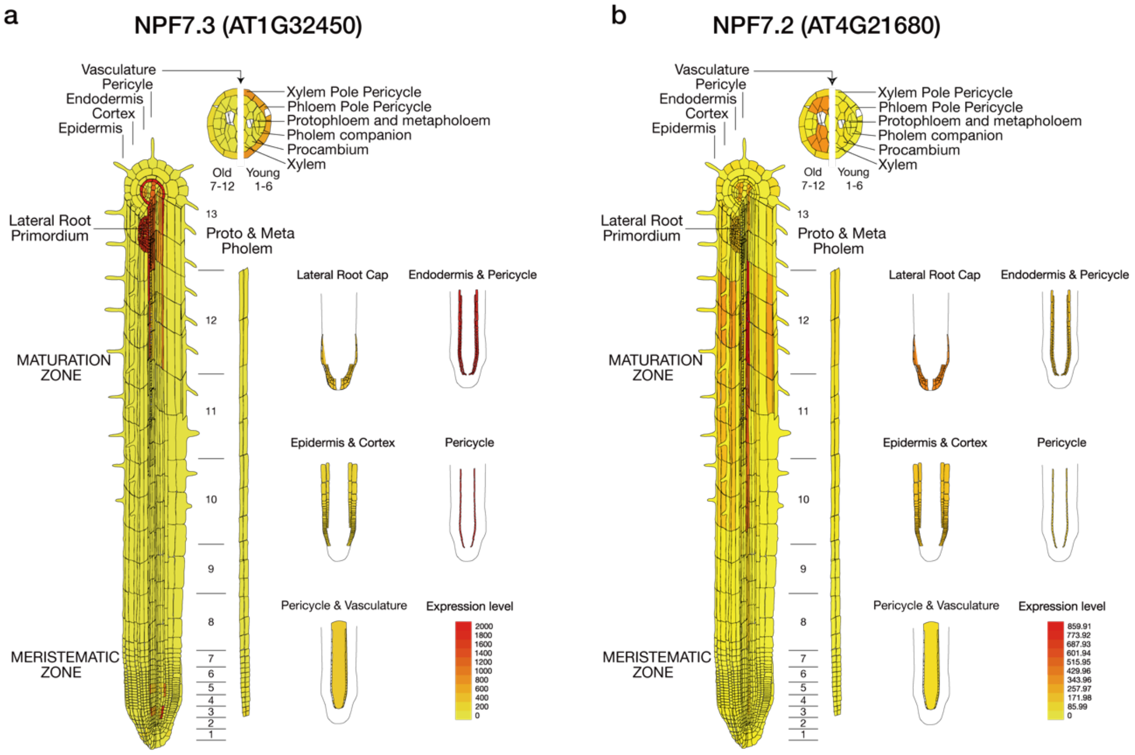


**Supplementary Fig. 4. Expression pattern of *NPF7.3*** Spatial expression patterns of *NPF7.3* (**a**) and *NPF7.2* (**b**) obtained from the Arabidopsis eFP Browser (http://bar.utoronto.ca/efp/cgi-bin/efpWeb.cgi)[1-3] are shown.

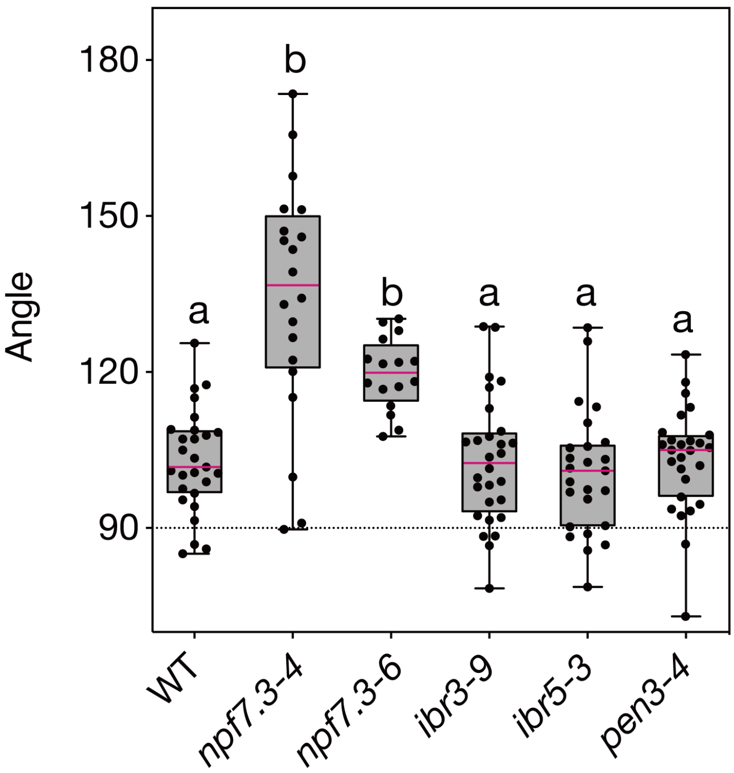


**Supplementary Fig. 5. Root gravitropism of IBA-related mutants.** Gravitropic responses of mutants defective in IBA-to-IAA conversion (*ibr3-9*), responses to auxin (*ibr5-3*) and IBA export (*pen3-4*) were analyzed. Vertical plates containing seven-day-old seedlings were rotated 90º, and root angles were measured one day after plate rotation. Dots represent individual measurements, and whiskers represent the entire spread of the data. Different letters indicate statistically significant differences by Dunn's multiple comparison test ($P < 0.05$).

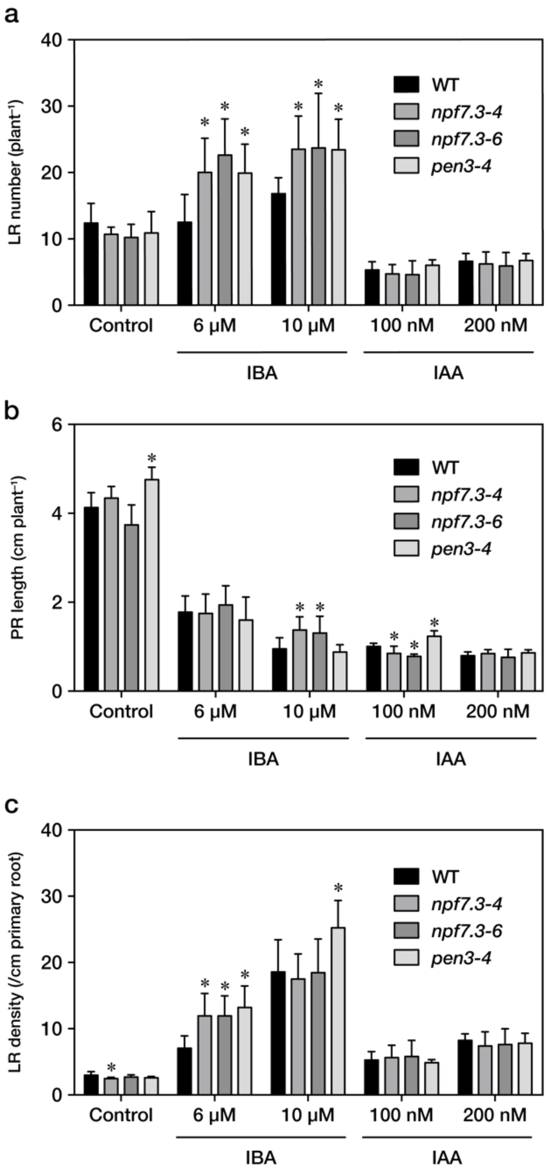


**Supplementary Fig. 6. Effects of IBA and IAA on lateral root formation and primary root elongation in *npf7.3*. a-c.** lateral root (LR) numbers (**a**), primary root (PR) lengths (**b**) and LR densities (**c**) of wild type (WT), *npf7.3* (*npf7.3-4* and *npf7.3-6*) and *pen3-4* treated with IBA or IAA. Two days after germination on half-strength MS media, seedlings were transferred to media containing IBA or IAA at the indicated concentrations. After further incubation for five days, LR numbers and PR lengths were measured. LR densities were calculated as the LR numbers divided by the PR lengths. Asterisks indicate significant differences.

### Supplementary Table 1. Primers used in this study.

| Target* | Use for | Forward primer | | Reverse primer | |
|---|---|---|---|---|---|
| | | Name | Sequence (5'-3') | Name | Sequence (5'-3') |
| ***npf7.3* mutant isolation** | | | | | |
| NPF7.3/NRT1.5 | PCR genotyping | NPF7.3g- F1 | CACGTTCACGTAAAATTCGAAG | NPF7.3g- R1 | AGCATGATCAAAAGGACATGC |
| NPF7.3/NRT1.5 | | NPF7.3g- F2 | AAAACGCGAGTTAGGAAGAGC | NPF7.3g- R2 | TAGAAAAACCCCAACCACATG |
| T-DNA | | LBb1.3 | ATTTTGCCGATTTCGGAAC | | |
| T-DNA | | O8409 | ATATTGACCATCATACTCATTGC | | |
| **Complementation test** | | | | | |
| NPF7.3 promoter | Construction of pENTR-NPF7.3pro | NPF7.3pro-topo-F | CACCCATCCCGTATGAGTATCAACCCAAAC | NPF7.3pro-R | TTTGCGATGATATATGATTATATGTATGTGAG |
| NPF7.3 promoter | Construction of pDONR-NPF7.3pro:NPF | NPF7.3pro-attL1-IF-F | GCCGCCCCCTTCACCCATCCCGTATGAGTATCAAC | AtNPF7.3pro-NPF7.3CDS-IF-R | CTCTAGGCAAGACATTTTGCGATGATATATGATTA |
| pDONR207-NPF7.3 CDS | 7.3 | NPF7.3CDS-IF-F | ATGTCTTGCCTAGAGATTTA | pDONR-attL1-R | GGTGAAGGGGGCGGCCGCGGAGCCTGCTTTTTTGTA |
| NPF7.3 promoter | Construction of pENTR-NPF7.3pro:NPF | NPF7.3pro-attL1-IF-F | GCCGCCCCCTTCACCCATCCCGTATGAGTATCAAC | AtNPF7.3pro-NPF7.2CDS-IF-R | AACTTTTTGATCCATTTTGCGATGATATATGATTA |
| pENTR-NPF7.2 CDS | 7.2 | NPF7.2CDS-IF-F | ATGGATCAAAAAGTTAGACA | pENTR-attL1-R | GGTGAAGGGGGCGGCCGC |
| NPF7.3 promoter | Construction of pENTR-NPF7.3pro:NPF | NPF7.3pro-attL1-IF-F | GCCGCCCCCTTCACCCATCCCGTATGAGTATCAAC | AtNPF7.3pro-NPF4.6CDS-IF-R | TTCTTCCACTTCCATTTTGCGATGATATATGATTA |
| pENTR-NPF4.6 CDS | 4.6 | NPF4.6CDS-IF-Fp | ATGGAA GTGGAAGAAG AGGT | pENTR-attL1-R | GGTGAAGGGGGCGGCCGC |
| **NPF7.3 promoter-reporter analysis** | | | | | |
| NPF7.3 promoter | Construction of pENTR-NPF7.3pro:GUS | NPF7.3pro-IF-F | GCCCCCTTCACCATGCATCCCGTATGAGTATCAAC | NPF7.3pro-GUS-IF-R | TACAGGACGTAACATTTTGCGATGATATATGATTATAT |
| GUS | | GUS-F | ATGTTACGTCCTGTAGAAAC | GUS-IF-R | AGCTGGGTCGGCGCGCTCATTGTTTGCCTCCCTGC |
| pENTR | | pENTR-F | AAGGGTGGGCGCGCCGACC | pENTR-R | GGTGAAGGGGGCGGCCGCG |
| NLS | Construction of | GFP-NLS-IF-F | CTGTACAAGTCCGGACTCAGATCTCGAG | GFP-NLS-IF-R | CTGGGTCTAGATATCTCGATTATCTAGATCCG |
| GFP | pENTR4-GFP-NLS | GFP-IF-F | AAAAGCAGGCTCCACCATGATGGTGAGCAAGGGCGAG | GFP-IF-R | TGAGTCCGGACTTGTACAGCTCGTCCATGC |
| GFP-NLS | Construction of pENTR4-NPF7.3pro:GFP-NLS | NRT7.3pro-GFP-NLS-IF-F | ATAATCATATATCATCGCAAAATGGTGAGCAAGGGCGAG | NRT7.3pro-GFP-NLS-IF-R | AGAAAGCTGGGTCGGCGCGTTATCTAGATCCGGTGAATCCTACC |
| pENTR-NPF7.3pro | Construction of pENTR4-NPF7.3pro:GFP-NLS | pENTR-NPF7.3pro-F | CGCGCCGACCCAGCTTTCTTGTACA | pENTR-NPF7.3pro-R | TTTGCGATGATATATGATTATATGTATGTGAG |

*Gene symbol provided by TAIR (http://www.arabidopsis.org/) except for T-DNA.

**Supplementary Table 2. Conditions of LC.**

| Solvent A | Solvent B | Gradient (composition of solvent B) | Column |
|---|---|---|---|
| Water containing 0.01% (v/v) acetic acid | MeCN containing 0.05% (v/v) acetic acid | Constant at 20% for 1 min | ZORBAX Eclipse XDB-C18 column, 2.1 x 50 mm i.d., 18 µm (Agilent) |
| | | Linear gradient from 20 to 35% over 4 min | |
| | | Linear gradient from 35 to 98% over 0.1 min | |
| | | Constant at 98% for 0.9 min | |
| | | Linear gradient from 98 to 20% over 0.1 min | |
| | | Constant at 20% for 0.9 min | |

**Supplementary Table 3. Parameters of tandem mass spectrometer.**

| Compound | Material | Retention time on LC (min) | Polarity of ESI | IonSpray voltage (kV) | Desolvation temperature (°C) | Declustering potential (V) | Collision energy (V) | Precursor ion (*m*/*z*) | Scan range (*m*/*z*) | Qualifier ion (*m*/*z*) |
|---|---|---|---|---|---|---|---|---|---|---|
| [$^{13}C_8$, $^{15}N_1$]IBA | Arabidopsis | 4.0 | + | 5.5 | 600 | 90 | 20 | 213.15 | 50-1000 | 195 |
| IBA | | 4.0 | + | 5.5 | 600 | 90 | 20 | 204.15 | 50-1000 | 186 |
| [methylene-$^2H_2$]IAA | | 2.5 | + | 5.5 | 600 | 90 | 15 | 178.20 | 50-1000 | 132 |
| IAA | | 2.5 | + | 5.5 | 600 | 90 | 15 | 176.20 | 50-1000 | 130 |
| [methylene-$^2H_2$]IBA | Yeast | 4.0 | – | –3.5 | 600 | –90 | –22 | 204.15 | 50-1000 | 160 |
| IBA | | 4.0 | – | –3.5 | 600 | –90 | –22 | 202.15 | 50-1000 | 158 |
| [methylene-$^2H_2$]IAA | | 2.5 | – | –3.5 | 600 | –90 | –15 | 176.20 | 50-250 | 132 |
| IAA | | 2.5 | – | –3.5 | 600 | –90 | –15 | 174.20 | 50-250 | 130 |
